# SMN1 and SMN2 Copy Number Determination by droplet digital PCR (ddPCR) based on Extreme Value Theory for Threshold Estimation

**DOI:** 10.1101/220020

**Authors:** Mohammad Niknazar, Michael Jansen, Oscar Puig

## Abstract

Spinal muscular atrophy (SMA) is the most frequent genetic cause of infantile death. Homozygous deletion screening of survival of motor neuron (SMN1) represents the first tier in diagnostic testing. In this work, we adopted and optimized a method to increase the accuracy of SMN1 and SMN2 copy number determination on ddPCR platform. This method, which makes no assumption about the distribution of the fluorescence readouts, was shown to significantly increase accuracy and outperform QuantaSoft software on problematic SMN1 samples.

**Method Summary:** In order to increase the accuracy of SMN1 and SMN2 copy number determination by ddPCR, we adopted and optimized a method based on extreme value theory for threshold estimation. Our method increases total accuracy and improves resolution of problematic samples.

## Main Body

Spinal muscular atrophy (SMA), an autosomal recessive neuromuscular disorder characterized by progressive loss of motor neurons in the spinal cord 1, is the most common cause of early childhood death (2,3). The homozygous deletion or mutation in the survival of motor neuron (SMN1) gene has been observed in the majority of SMA patients 4, but all patients retain a centromeric copy of the gene, SMN2 5. Since SMA is a severe disease with a high carrier frequency, direct carrier dosage testing has been beneficial to many families 6. Quantitative methods of SMN dosage use real-time PCR 7, but many factors can impact PCR efficiency and hence the threshold cycle, affecting accuracy and precision 8. Currently, the droplet digital PCR system of Bio-Rad is a widely used ddPCR platform. However, the threshold setting method in QuantaSoft software (Bio-Rad Laboratories, Inc., Hercules, CA, USA) that comes with the ddPCR platform is not always accurate as the threshold often falls within the population of negative droplets or is not possible to calculate. This leads to inaccuracies in copy number determination for SMN1 and SMN2. In this paper, we adopted and optimized the method described in 9 to estimate the threshold which is needed to determine copy numbers of SMN1 and SMN2. The method does not make any assumptions about the distribution of the fluorescence readouts and estimates the threshold by modelling the extreme values in the negative droplet population using extreme value theory. Moreover, this method takes into account shifts in baseline fluorescence between samples.

Clinical samples from a commercial carrier screening platform (CarrierMap, Recombine, Inc.) were used after previously failing SMA testing in an internal qPCR assay 10 and commercial ddPCR assay (Bio-Rad catalog number 1863500), respectively. These samples were sent to a CAP/CLIA laboratory (Athena Diagnostics Inc., Marlborough, MA, USA) for unequivocal SMN1 and SMN2 analysis. If a sample failed the ddPCR assay for any reason (<10,000 droplets, <100 copies/uL, too much “rain” to create a confident threshold, or a copy number outside of 0.2 copies from a whole number), it was noted as a failure and was not assigned a copy number. Some samples failed multiple times in the ddPCR assay with inconsistent results, were labeled as “problematic” samples and no copy number was obtained from ddPCR testing. Sample data was then exported from the QuantaSoft software to perform the analysis method discussed in this paper. A previously presented method 9, which makes no assumptions about the distribution of the fluorescence readouts was adopted and optimized to determine copy numbers of SMN1 and SMN2. This method relies on application of generalized extreme value (GEV) distribution for estimating a threshold that segregates positive and negative droplets in ddPCR systems. The raw data in no template controls (NTCs) are first normalized to correct for variations among NTCs. Robust mode estimator suggested by Robertson and Cryer 11 is applied to each NTC to obtain a stable estimate of measure of central tendency, which does not require any parameters to be set and is not dependent on the amount of rain. After subtracting the estimated robust mode from all measured intensities of each NTC, all the NTCs are merged to represent a single sample from the population of negative droplets 9. This normalization procedure by subtracting robust mode is applied even if there is only one NTC available.

To fit GEV distribution to fluorescence intensity values, the sample is divided to *β* blocks of subsamples and the distribution parameters are calculated by fitting to the maxima of the blocks. These parameters and an initial threshold, *T*, predetermined percentile of the fitted distribution, are used to obtain the threshold relative to normalized NTCs. We then need a raw threshold to classify negative and positive droplets, which can be obtained by averaging the robust modes of the NTCs and adding the threshold related to the NTCs. Then, after estimating the robust mode, it will be added to the previously calculated threshold (related to the NTCs) to obtain the final threshold of the sample.

Using the final threshold, positive and negative droplets can be classified and the concentration, *C*, is calculated as 9:

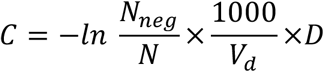

where V_d_ is 0.91 μL and D is dilution factor as described in 8. Finally, copy number is calculated as the ratio of the target molecule concentration to the reference molecule concentration, multiplied by the number of copies of reference species in the genome:

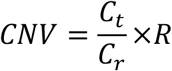

where *C_t_* is concentration of target species, *C_r_* is concentration of reference species, and *R* is a ratio which is considered as the number of copies of reference loci in the genome and is usually set as 2. However, since *C_t_* and *C_r_* are not necessarily obtained at the same speed and other identical conditions, we considered this value as a parameter to be estimated.

This method requires appropriate values for number of blocks *B*, initial threshold *T*, and the ratio *R*. In order to estimate these parameters for SMN1 and SMN2, we used validation samples (384 SMN1 and 288 SMN2) measured in an external CAP/CLIA laboratory. The parameters were estimated as *B* = 252 blocks per NTC, *T* = 0.99851459 and *R* = 1.7911230 for SMN1, and *B* = 256 blocks per NTC, T = 0.89866313 and R = 1.85829011 for SMN2.

After optimization, our method was applied to independent test samples. QuantaSoft software and our method provided similar accuracies in non-problematic samples (96 SMN1 and 130 SMN2): 95.70% (95% CI [91.64%, 99.76%]) and 97.77% (95% CI [94.82%, 100%]) for SMN1, and 86.15% (95% CI [80.21%, 92.09%]) and 86.92% (95% CI [81.12%, 92.72%]) for SMN2, respectively. However, our method provided significantly improved accuracy for 165 problematic SMN1 samples: 45.45% (95% CI [37.85%, 53.05%]) for QuantaSoft and 85.45% (95% CI [80.07%, 90.83%]) for our method. Quantasoft accuracy in a set of 12 samples processed several times with inconsistent results was 50.00% (95% CI [21.71%, 78.29%]), in contrast to 83.33% (95% CI [62.24%, 100%]) with our method.

In this work, we adopted ddpcRquant method which estimates the threshold by modeling the extreme values in the negative droplet population using extreme value theory while no assumptions are made about the distribution of the fluorescence readouts. We then optimized the parameter sets for SMN1 and SMN2 separately to determine their copy numbers. The results show that this method significantly outperforms QuantaSoft software in determining the copy numbers of SMN1 problematic/flagged samples. Future work is combining this method with other technologies to further increase the accuracy of copy number determination.

## Author contributions

M.N. optimized the method, analyzed the data, and wrote the paper. M.J. ran the samples and acquired the data. O.P. designed and led the research.

## References

1. Lunn M.R., C.H Wang. 2008. Spinal muscular atrophy. The Lancet. 371(9630):2120–33.

2. Ogino S., R.B. Wilson. 2002. Genetic testing and risk assessment for spinal muscular atrophy (SMA). Human genetics. 111(6):477–500.

3. Prior T.W., P.J. Snyder, B.D. Rink, D.K. Pearl, R.E. Pyatt, D.C. Mihal, T. Conlan, B. Schmalz, L. Montgomery, K. Ziegler, C. Noonan. 2010. Newborn and carrier screening for spinal muscular atrophy. American Journal of Medical Genetics Part A. 152(7):1608–16.

4. Kolb S.J., J.T. Kissel. 2011. Spinal muscular atrophy: a timely review. Archives of neurology. 68(8):979–84.

5. D‘Amico A., E. Mercuri, F.D. Tiziano, E. Bertini. 2011. Spinal muscular atrophy. Orphanet journal of rare diseases. 6(1):1.

6. Prior T.W. 2008. Carrier screening for spinal muscular atrophy. Genetics in Medicine. 10(11):840–2.

7. Feldkötter M., V. Schwarzer, R. Wirth, T.F. Wienker, B. Wirth. 2002. Quantitative analyses of SMN1 and SMN2 based on real–time lightCycler PCR: fast and highly reliable carrier testing and prediction of severity of spinal muscular atrophy. The American Journal of Human Genetics. 70(2):358–68.

8. Pinheiro L.B., V.A. Coleman, C.M. Hindson, J. Herrmann, B.J. Hindson, S. Bhat, K.R. Emslie. 2011. Evaluation of a droplet digital polymerase chain reaction format for DNA copy number quantification. Analytical chemistry. 84(2):1003–11.

9. Trypsteen W., M. Vynck, J. De Neve, P. Bonczkowski, M. Kiselinova, E. Malatinkova, K. Vervisch, O. Thas, L. Vandekerckhove, W. De Spiegelaere. 2015. ddpcRquant: threshold determination for single channel droplet digital PCR experiments. Analytical and bioanalytical chemistry. 407(19):5827–34.

10. Maranda, B., L. Fan, J.F. Soucy, L. Simard, G.A. Mitchell. 2012. Spinal muscular atrophy: Clinical validation of a single–tube multiplex real time PCR assay for determination of SMN1 and SMN2 copy numbers. Clinical biochemistry, 45(1), pp.88–91.

11. Robertson T., J.D. Cryer. 1974. An iterative procedure for estimating the mode. Journal of the American Statistical Association. 69(348):1012–6.

